# IgSeqR: a protocol for the identification, assembly, and characterization of full-length tumor Immunoglobulin transcripts from unselected RNA sequencing data

**DOI:** 10.1101/2024.09.03.611002

**Authors:** Dean Bryant, Benjamin Sale, Giorgia Chiodin, Dylan Tatterton, Benjamin Stevens, Alyssa Adlaon, Erin Snook, James Batchelor, Alberto Orfao, Francesco Forconi

## Abstract

Immunoglobulin (IG) gene analysis provides fundamental insight into B-cell receptor structure and function. In B-cell tumors, it can inform the cell of origin and clinical outcomes. Its clinical value has been established in the two types of chronic lymphocytic leukemia with unmutated or mutated *IGHV* genes and is emerging in other B-cell tumors. The traditional PCR-based techniques, which are labor-intensive, rely on the attainment of either a dominant sequence or a small number of subclonal sequences and do not allow automated matching with the clonal phenotypic features. Extraction of the expressed tumor IG transcripts using high-throughput RNA sequencing (RNA-seq) can be faster and allow the collection of multiple sequences matched with the transcriptome profile. Analytical tools are regularly sought to increase the accuracy, depth, and speed of acquisition of the full *IGV-(IGD)-IGJ-IGC* sequences and combine the *IG* characteristics with other RNA-seq data. We provide here a user-friendly protocol for the rapid extraction, identification, and accurate determination of the full (leader to constant region) tumor *IG* templated and non-templated transcript sequence from RNA-seq. The derived amino acid sequences can be interrogated for their physico-chemical characteristics and, in certain lymphomas, predict tumor glycan types occupying acquired N-glycosylation sites. These features will then be available for association studies with the tumor transcriptome. The resulting information can also help refine diagnosis, prognosis, and potential therapeutic targeting in the most common lymphomas.

## Introduction

The B-cell receptor (BCR) immunoglobulin (IG) glycoprotein is the defining functional feature of a mature B cell, and *IG* gene analysis can provide fundamental insight into the origin and behavior of a B-cell tumor [1, 2]. It is a Y-shaped dimer of 2 identical heavy and light chains, with 2 main functional components. A variable region that confers diversity to recognize different antigens and is unique to each B cell, and a constant region with an effector function. *IG* diversity results from a series of genetic recombinations at the *IG* heavy (*IGH*) and kappa (*IGK*) or lambda (*IGL)* light chain loci during B cell development in the bone marrow before a naïve B cell exits to the periphery (**Figure 1**). For the heavy chain, the recombinations are accompanied by non-templated nucleotide additions/deletions at the junctions of one of ∼51 *IGHV*, ∼21 *IGHD*, and ∼6 *IGHJ* genes in the complementarity-determining region 3 (*CDR3*) forming the “fingerprint” of an individual B cell. Further variability is conferred by the recombination of a *V* gene with *a J* gene at the *IGK* or *IGL* loci. Following antigen encounter, naïve B-cells undergo class-switch recombination and somatic hypermutation, typically in the presence of activation-induced cytidine deaminase (AID), T cells, and cytokines, for affinity maturation in the germinal center (GC) and differentiation in memory B cells or plasma cells [3]. The GC reaction involves proliferation, which makes the B cells vulnerable to damage and transformation into tumors. Tumor B cells preserve the *IG* sequence of the cell of origin. Therefore, analysis of the *IG* sequences allows the identification of the stage of differentiation reached by a B-cell before tumor transformation [4-6].

**Figure 1.**
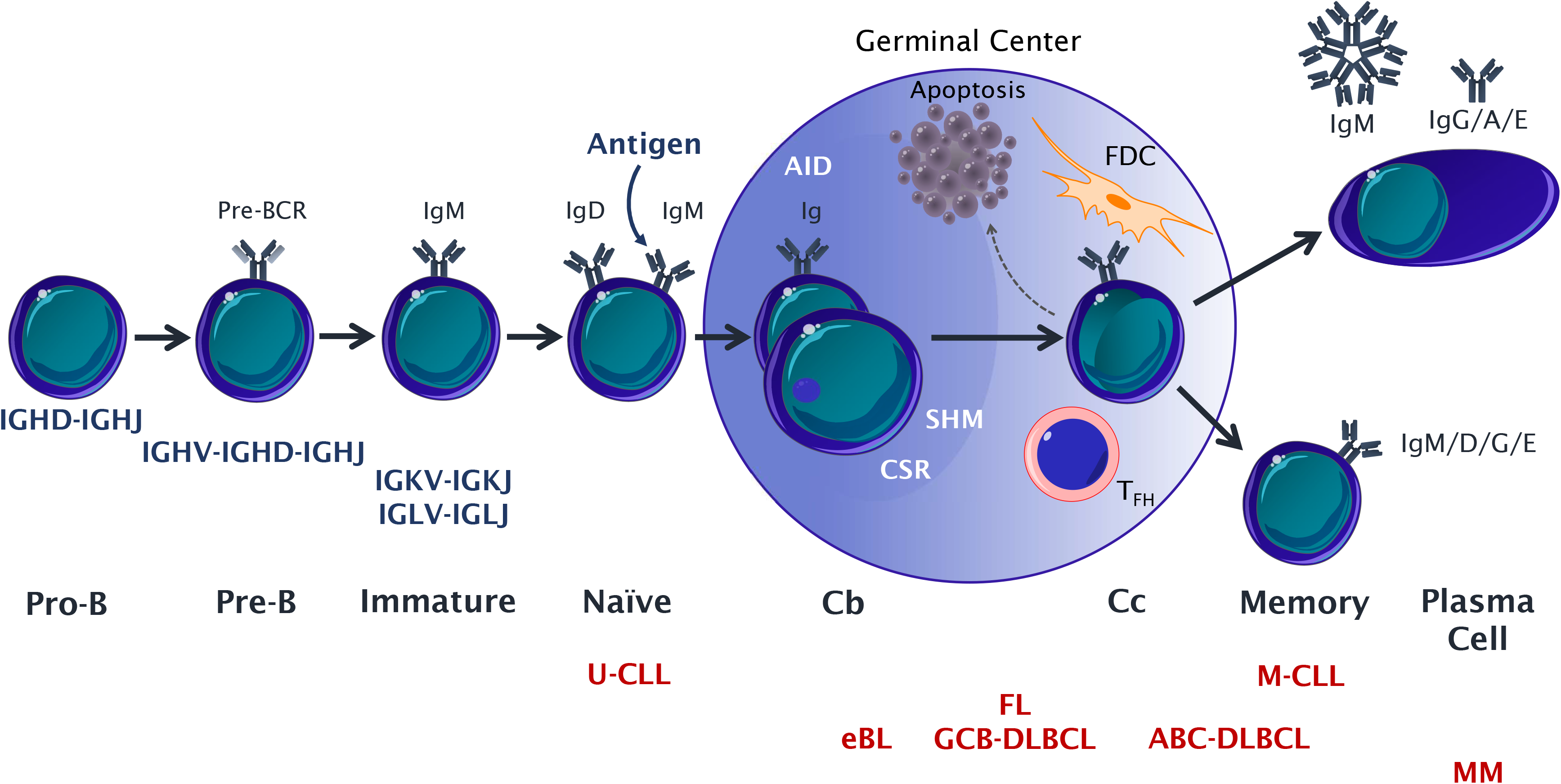
Immunoglobulin gene analysis provides insight into the cell of origin and behavior of B-Cell malignancies. The immunoglobulin heavy-chain gene repertoire comprises ∼51 functional variable (*IGHV*), ∼21 diversity (*IGHD*), and ∼7 joining (*IGHJ*) genes at the *14q32* locus. In the bone marrow, progenitor B cells (Pro-B cells) undergo an *IGHD-IGHJ* rearrangement. If successful, a complete *IGHV-IGHD-IGHJ* rearrangement occurs at the precursor B cell (Pre-B), which expresses a precursor B-cell receptor (pre-BCR) containing a surrogate VpreB1 light chain and a full IG heavy chain. The pre-BCR will promote rearrangement of *IGKV-IGKJ* at the *2p11*.*2* locus, and, if this is non-functional in both alleles, the rearrangement of *IGLV-IGLJ* will occur at the *22q11*.*22* locus in immature B cells. A successful rearrangement of the IG light chain enables the expression of a competent immunoglobulin M (IgM) and autoreactive B cell clones within the bone marrow microenvironment will be deleted, ensuring the production of functional non-autoreactive naïve B cells expressing IgM and IgD. IgM+ve IgD+ve naïve B cells exit the bone marrow and migrate to peripheral lymphoid organs (spleen, lymph nodes, MALTs, etc.) where they will encounter antigen, and they will undergo class-switch recombination (CSR) and somatic hypermutation (SHM) in the presence of activation-induced cytidine-deaminase (AID) in a germinal center (GC) reaction at the centroblast (Cb) stage (dark zone). During SHM, Cb introduce point mutations in the IG variable region genes to mature affinity to antigen. Centrocytes (Cc) emerge in the light zone where their fate will depend on their BCR interactions with immune complexes on follicular dendritic cells (FDC) in the presence of T follicular helper (T_FH_) cells. Cc with the BCR of the right affinity to antigen receive survival signals and differentiate into memory B cells or plasma cells, while the others will undergo apoptosis. The tumor *IG* genes preserve the features of the cell having undergone transformation. Chronic Lymphocytic Leukemias with unmutated *IG* genes (U-CLL) arise from pre-GC B-cells and have an aggressive clinical course, while those with mutated *IG* genes (M-CLL) arise from post-GC B cells and display an indolent clinical course. In endemic Burkitt lymphoma (eBL), FL, and some DLBCL, there is intraclonal heterogeneity of the *IGV* gene sequences to indicate that the SHM process is ongoing, as in a GC B cell. Diffuse Large B-cell Lymphoma (DLBCL) can be classified into two major subtypes: GC B-cell-like (GCB) and activated B-cell-like (ABC). Asparagine-x-serine/threonine N-glycosylation motifs (where X is any amino acid except proline) are introduced by SHM, allowing occupation of the sites by oligomannose-type glycans in almost all FL and in ∼30% of all GCB-DLBCL. Multiple myeloma (MM) is characterized by the clonal expansion of plasma cells, which carry mutated *IG* and secrete a monoclonal IG in the serum (paraprotein).

In chronic lymphocytic leukemia (CLL), *IG* analysis reveals two major types defined by *IGHV* mutational status [5]. The CLL type with unmutated *IGHV* (U-CLL) derives from pre-germinal center CD5^+^ B cells, while the CLL type with mutated *IGHV* (M-CLL) appears to arise from post-follicular CD5^+^ B cells [7, 8]. Since the discovery that U-CLL has a worse prognosis than M-CLL [9, 10], subsequent studies have demonstrated that each type has a distinctive cellular origin, biology, (epi)genetics, clinical prognosis, and response to therapy [5, 11]. *IGHV* gene analysis has become an essential part of the diagnostic workup for any patient with CLL.

*IG* analysis also informs key tumor-specific features in certain lymphomas. In classic follicular lymphoma (FL), the tumor *IG* acquires N-glycosylation sites (AGS), defined by the asparagine-X-serine/threonine motif (where X is any amino acid except proline) [12]. AGS in FL are typically in the CDRs of the variable region by somatic hypermutation [13] and are occupied by tumor-specific oligomannose-type glycans [14-16]. These atypical glycans are unique to the tumor B cell, are present on the entire FL clone, and persist during the entire clonal history of FL through transformation into diffuse large B-cell lymphoma (DLBCL), despite ongoing somatic hypermutation [14, 17].

The current gold standard for *IG* gene analysis is by Sanger sequencing. This approach offers a highly accurate *IG* sequence but is time-consuming, labor-intensive, and requires a dedicated experimental and analytical workflow on samples with documented high tumor infiltration [18]. The increasing adoption of high-throughput whole transcriptome RNA sequencing (RNA-seq) methods allows many tests to be streamlined into a single experimental workflow. Through the application of appropriate analytical pipelines, a single RNA-seq experiment can yield comprehensive information on gene expression, isoform expression, single nucleotide polymorphisms (SNPs), and larger structural variants [19].

RNA-seq can therefore be a better alternative to Sanger sequencing in *IG* gene analysis. However, the intrinsic high variability of the non-templated CDR3 sequences has been a challenge to the identification of the full *IG* sequence with the current RNA-seq analytical workflows, which have involved mapping reads to a reference transcriptome.

Here we describe IgSeqR (pronounced I-G-Seeker), a protocol for the reference-free extraction, identification, and accurate determination of the full tumor *IG* transcript sequence from unselected whole transcriptome RNA-seq data.

### Development of the protocol

We first used IgSeqR to identify the tumor *IG* full transcripts in a cohort of 489 DLBCL with RNA-seq data publically available [14]. The data were deposited in the National Cancer Institute (NCI) Genomic Data Commons (accession phs001444.v1.p1) [20, 21]. The full *IGHV*-*IGHD*-*IGHJ* sequence rearrangements were identified from leader to constant region with high confidence in 339 (69%) samples, from which we could determine *IGHV, IGHD, IGHJ*, and *IGHC* use, homology to germline, and AGS presence and location. Since we were interested in those cases with N-glycosylation sites acquired by somatic hypermutation and no information was available on the tumor purity of these samples, we investigated only the 307 samples with mutated (<98% homology to germline) *IGHV* [14]. We found that the AGS were preferentially in the EZB genetic subtype of the GC-B-cell-like (GCB) DLBCL. The majority of these AGS were located in the CDR, in a fashion similar to FL. Following the generation of F(ab) from the tumor-derived *IG* heavy and light chain sequences we documented that the glycan structure occupying the AGS was location-dependent and that the oligomannose-type glycans occupied the CDR-located sites only. We performed correlation studies with the transcriptome profile and defined genes and gene sets differentially expressed in samples with and without AGS. We performed correlations with the clinical characteristics of the DLBCL. Interestingly, we found that AGS in the EZB subtype conferred a poor prognosis, indicating that this approach for *IG* gene analysis could be adopted to predict both glycan structure and response to conventional therapies [14]. In the present study, we report the IgSeqR script while validating its accuracy in primary CLL samples with matched *IG* heavy chain Sanger and bulk RNA-seq data (deposited in ArrayExpress, accession E-MTAB-12017) [22]. IgSeqR is fully concordant with Sanger sequencing for *IGHV, IGHD*, and *IGHJ* allele use and nucleotide sequence.

### Applications of the method

IgSeqR is ideal for studies requiring high-quality base calls across the full sequence, including the non-templated CDR3 region, of the *IG* heavy and light chains of any mature B cell tumor. The protocol reduces the computational burden of *de novo* assembly by pre-filtering redundant data and allows the identification of the dominant nucleotide sequence of the *IG* heavy and light chains from leader to constant region from RNA-seq data. Through the alignment to the most updated *IG* sequence repertoires, currently IMGT/V-QUEST reference directory 202349-3, program version 3.6.2 at http://www.imgt.org, it is possible to obtain insights into *IGHV, IGHD, IGHJ* heavy chain alleles, *IGKV, IGKJ* or *IGLV, IGLJ* light chain alleles, constant region class and subclass, homology to germline, CDR1-3 and FR1-4 characteristics.

IgSeqR can also be applied to autoimmune and infectious diseases to identify common patterns recurring in the polyclonal expansions (e.g. dominance and characteristics of *IGHV1-69* in rheumatoid arthritis or influenza) [23, 24].

IgSeqR can also be used to generate F(ab)s [14] or improve strategies for vaccine and antibody therapy development [25-27].

### Comparisons with other methods

Compared to Sanger sequencing and existing RNAseq-based protocols, IgSeqR increases the length of the transcript containing the full *IGV*(-*IGD*)-*IGJ* rearrangements from leader to *IG* constant region (up to 2000 nucleotides).

It maintains the same level of accuracy as Sanger sequencing while improving the chance of detecting a clonal sequence compared to a PCR-based approach, particularly in lymphoma samples. *IG* sequencing of lymphoma samples by Sanger is notoriously difficult and demands significant amounts of equipment and time, particularly if subcloning approaches are necessary, to identify small cohorts of patients [28-30]. In a cohort of 37 lymphomas with more than 10% tumor B cells in the test sample by flow cytometry, PCR/direct Sanger sequencing successfully identified a dominant *IG* rearrangement in only 11 (30%). By Cibersort estimation [31], 439 DLBCL samples from the NCI cohort had > 10% (tumor) B cells. IgSeqR identified the tumor *IG* rearrangement in 319 (73%), a significantly superior frequency than Sanger (p<0.0001). However, IgSeqR was also successful in identifying the full *IG* sequence in 20 of the 50 (40%) samples with <10% B cell purity, although the success rate was lower compared to >10% (p<0.005) (**Figure 2** and **Table S1**).

**Figure 2.**
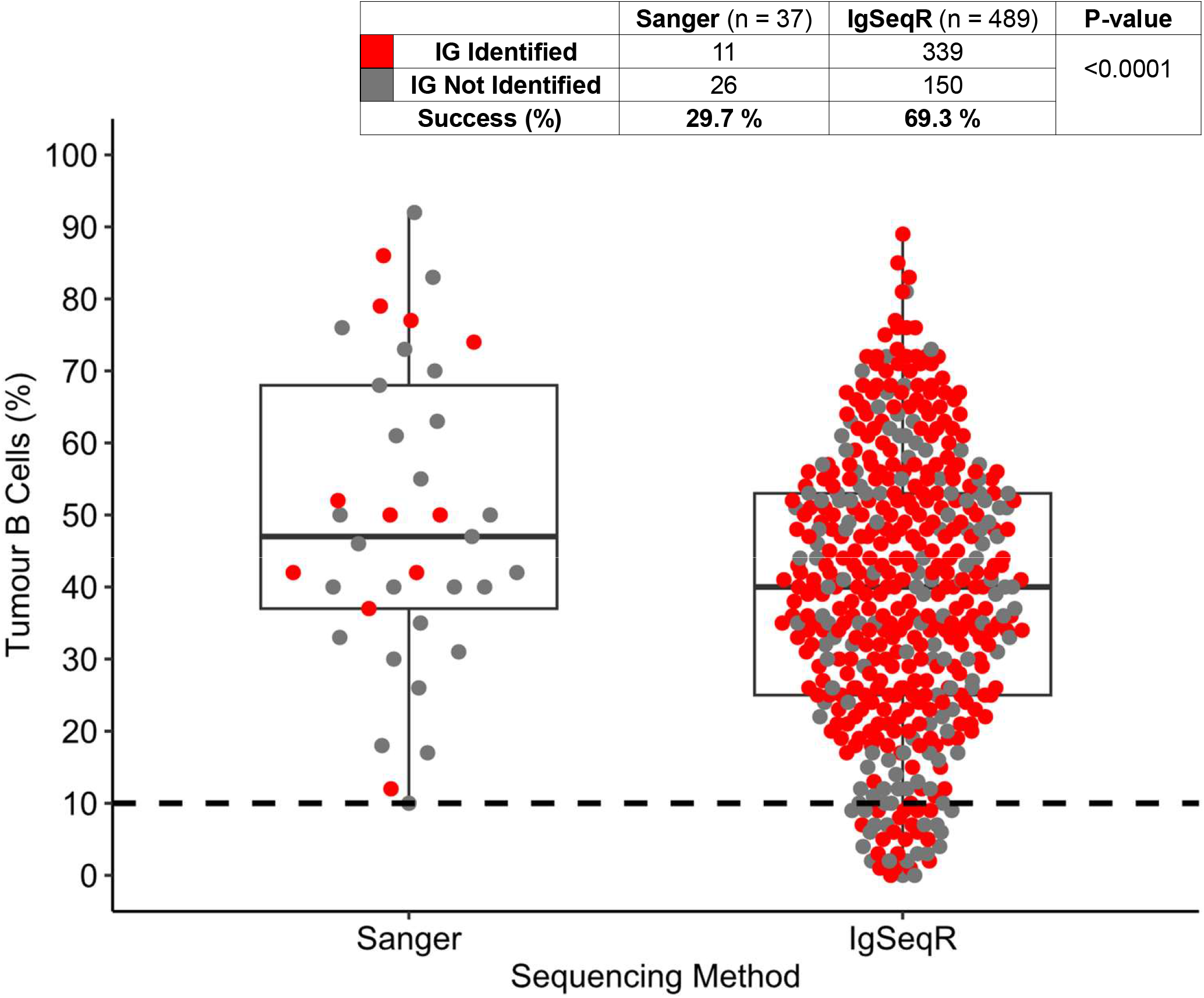
Tissue-derived lymphoma samples, in which the tumor *IG s*equence was sought by Sanger or IgSeqR. Each dot identifies a sample: red dots indicate the samples where the tumor *IG* sequence was identified; grey dots indicate the samples where the tumor *IG* sequence could not be identified. The accompanying table indicates the number and proportion of *IG* sequences identified using the individual methods. There was a significantly higher proportion and probability of identifying the tumor *IG* sequence by RNA-seq/IgSeqR (69%) compared to PCR/Sanger (30%) (X-square with Yates’ correction, p-value <0.0001). Sequencing by Sanger was performed only in samples with >10% tumor infiltration by immunophenotype, while IgSeqR was applied to any sample irrespective of (tumor) B cell percentage, as estimated by Cibersort.

The experimental and analytical time to identify the sequences by Sanger was in weeks, while it was in days for the IgSeqR approach. This suggests that IgSeqR is dramatically efficient, offering a higher success rate in a shorter experimental and analytical time compared to standard PCR and Sanger sequencing.

Although IgSeqR is currently not configured to build the *IGHC* sequence with contigs spanning from CDR3 to the 3’ end of the constant region allele used, the derived transcripts recovered are generally sufficient to determine the *IGHC* class and subclass with high confidence. This is another advantage compared to Sanger, where individual isotypes can only be identified using isotype-specific primers.

Several tools have been developed for *IG* analysis from bulk and single-cell RNA-seq [32-40] (**Table 1**), many of which preferentially rely on aligning RNA-seq reads to *IG* reference sequences [33, 35-37]. MiXCR is widely adopted for immune profiling in both academic and industrial settings [33]. It primarily uses the N-regions at the *IGV*-(*IGD*)-*IGJ* junctions as a reference and identifies and quantifies the *IG* repertoire by CDR3 diversity. However, it is less focused on the full length, and highly mutated *IGV*-(*IGD*)-*IGJ* sequences may not be fully reconstructed. TRUST4 and IG_ID tools utilize *de novo* transcriptome assembly. However, TRUST4 was initially designed for TCR, rather than BCR, repertoire analysis [39]. The IG_ID tool can accurately produce full-length BCR transcripts comparable to Sanger sequencing, but has an extended processing time and generates large temporary files due to the *de novo* assembly of the whole transcriptome, limiting its use for large-scale analyses [32].

**Table 1.** Published tools for IG analysis from bulk and single-cell RNA sequencing (RNA-seq)

| Tool | Description | Receptor | Sequencing Data | Reference |
| --- | --- | --- | --- | --- |
| IG_ID | *De Novo* assembly of BCR transcripts from bulk RNA-seq data | BCR | Bulk | Blachly et al 2015 [31] |
| MiXCR | Analysis of raw T- or B- cell receptor repertoire sequencing data | BCR/TCR | Bulk/Single Cell | Bolotin et al 2015 [32] |
| BASIC | Bayesian inference of immunoglobulin sequences. It offers functionalities for V(D)J gene identification, clonotype analysis, and mutation profiling. | BCR | Single Cell | Canzar et al 2017 [33] |
| IMSEQ | Provides functionalities for the identification and quantification of IG genes, as well as the detection of somatic hypermutations | BCR/TCR | Bulk | Kuchenbecker et al 2015 [34] |
| ImReP | Extraction of receptor reads from sequencing data and assemble clonotypes, detect corresponding V(D)J recombinations and correct PCR sequencing errors | BCR/TCR | Bulk | Mandric et al 2020 [35] |
| V'DJer | Customized read extraction, assembly and V(D)J rearrangement detection and filtering to produce contigs representing the most abundant portions of the BCR repertoire | BCR | Bulk | Mose et al 2016 [36] |
| VDJPuzzle | Provides a user-friendly interface for the identification of V(D)J rearrangements, clonotype analysis, and visualization of TCR and BCR repertoires. | BCR/TCR | Single Cell | Rizzetto et al 2018 [37] |
| TRUST4 | Performs de novo assembly on V, J, C genes including the hypervariable complementarity-determining region 3 (CDR3) and reports consensus of BCR/TCR sequences | BCR/TCR | Bulk/Single Cell | Song et al 2021 [38] |
| BALDR | Infers the clonal structure of B-cell repertoires, providing information on clonal abundance, V(D)J gene usage, and somatic hypermutations | BCR | Single Cell | Upadhyay et al 2018 [39] |

We performed a direct comparison of IgSeqR with the MiXCR (v 4.3.2) or TRUST4 (v1.0.12) with the 18 CLL samples (**Tables 2** and **S2**).

**Table 2.** A comparison between RNA-seq based Immunoglobulin Gene analysis tools, IgSeqR MiXCR and TRUST4.

| Property | IgSeqR | MixCR | TRUST4 |
| --- | --- | --- | --- |
| Recognized as IG (IMGT) | 18 (100 %) | 18 (100 %) | 17 (94.44 %) |
| Productive Sequence | 18 (100 %) | 18 (100 %) | 17 (94.44 %) |
| Complete VDJ | 18 (100 %) | 17 (94.44 %) | 17 (94.44 %) |
| IGHV Gene match | 18 (100 %) | 17 (94.44 %) | 17 (94.44 %) |
| IGHV Seq match | 18 (100 %) | 14 (77.78 %) | 17 (94.44 %) |
| IGHD Gene Allele match | 18 (100 %) | 18 (100 %) | 17 (94.44 %) |
| IGHD Seq match | 18 (100 %) | 18 (100 %) | 17 (94.44 %) |
| IGHJ Gene Allele match | 18 (100 %) | 18 (100 %) | 17 (94.44 %) |
| IGHJ Seq match | 18 (100 %) | 18 (100 %) | 17 (94.44 %) |
| CDR3 Seq match | 18 (100 %) | 18 (100 %) | 17 (94.44 %) |
| Full Sanger VDJ Concordance | 18 (100 %) | 14 (77.78 %) | 17 (94.44 %) |
| Average Length | 2036 | 589 | 768 |
| Assembly efficiency (Seconds/Nucleotide) | 1.18 | 8.10 | 1.44 |

In red are identified the properties of MixCR or TRUST4 with inferior performance compared to IgSeqR.

MiXCR generated *IGH* transcripts for all 18 of the samples, but only 17 (94%) of these spanned the full *IGHV*-*IGHD*-*IGHJ* rearrangement, and only 14 (78%) had 100% identity with Sanger.

TRUST4 generated *IGH* transcripts from 17 (94%) of the samples, all of which were fully concordant with Sanger. However, TRUST4 failed to identify the only case that had a deletion of codon 66 of the *IGHV4-34* tumor sequence, possibly revealing a limitation of TRUST4 in identifying insertions or deletions.

IgSeqR also produced the longest tumor transcripts, averaging a length of 2036 nucleotides, compared to 589 and 769 nucleotides by MiXCR and TRUST4 respectively. Notably, the majority (78%) of the IgSeqR transcripts were long enough to cover the full *IGH* region from leader to the membrane domain of the constant region with confidence, a feature not possible in the shorter transcripts generated by MiXCR or TRUST4 (**Figure 3**).

**Figure 3.**
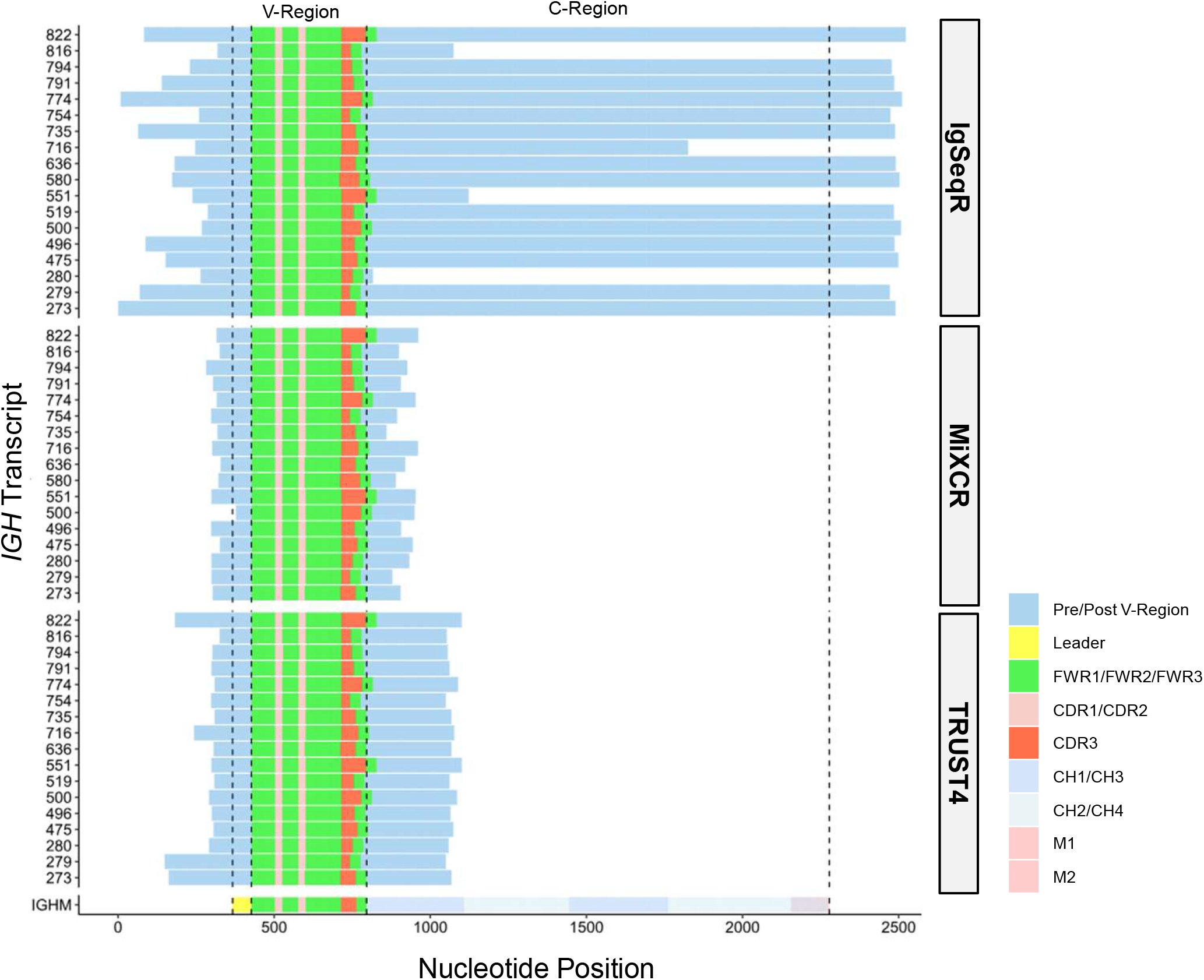
Comparison between the transcripts recovered by the IgSeqR pipeline with MiXCR, TRUST4 and a reference *IGHM* transcript. A direct comparison of three analytical tools for the recovery of *IGHV-IGHD-IGHJ* transcripts recovered from unselected bulk high throughput RNA sequencing data from 18 chronic lymphocytic leukemia samples with high tumor purity. The tools, IgSeqR, MiXCR (v 4.3.2), and TRUST4 (v1.0.12) were run using the Iridis5 high-performance computing cluster at the University of Southampton, utilizing 8 × 2.0 GHz CPU cores and 32 GB RAM to simulate a typical desktop workstation. The resulting transcripts were assessed for recovery of a full-length, productive *IGHV-IGHD-IGHJ* (V-Region) transcript and concordance with matched Sanger sequencing in the V-region. MiXCR recovered *IGHV-IGHD-IGHJ* transcripts for all 18 of the samples, with 17 (94%) having productive and full V-Region coverage, however only 17 used the same IGHV of Sanger, and 14 (78%) had 100% identity with Sanger. TRUST4 generated *IGHV-IGHD-IGHJ* transcripts from 17 (94%) of the samples, all of which had productive and full V-Region coverage and full concordance with Sanger. IgSeqR demonstrated productive and full V-Region coverage and full concordance with Sanger in all 18 (100%) samples. IgSeqR also produced the longest tumor transcripts, averaging a length of 2036 nucleotides, compared to 589 and 769 nucleotides by MiXCR and TRUST4 respectively. Notably, the majority (78%) of the IgSeqR transcripts were long enough to cover the full *IGHM* transcript from leader to the membrane domains (M1 and M2) of the constant region (C-Region), a feature not possible in the shorter transcripts generated by MiXCR or TRUST4.

**Figure 4.**
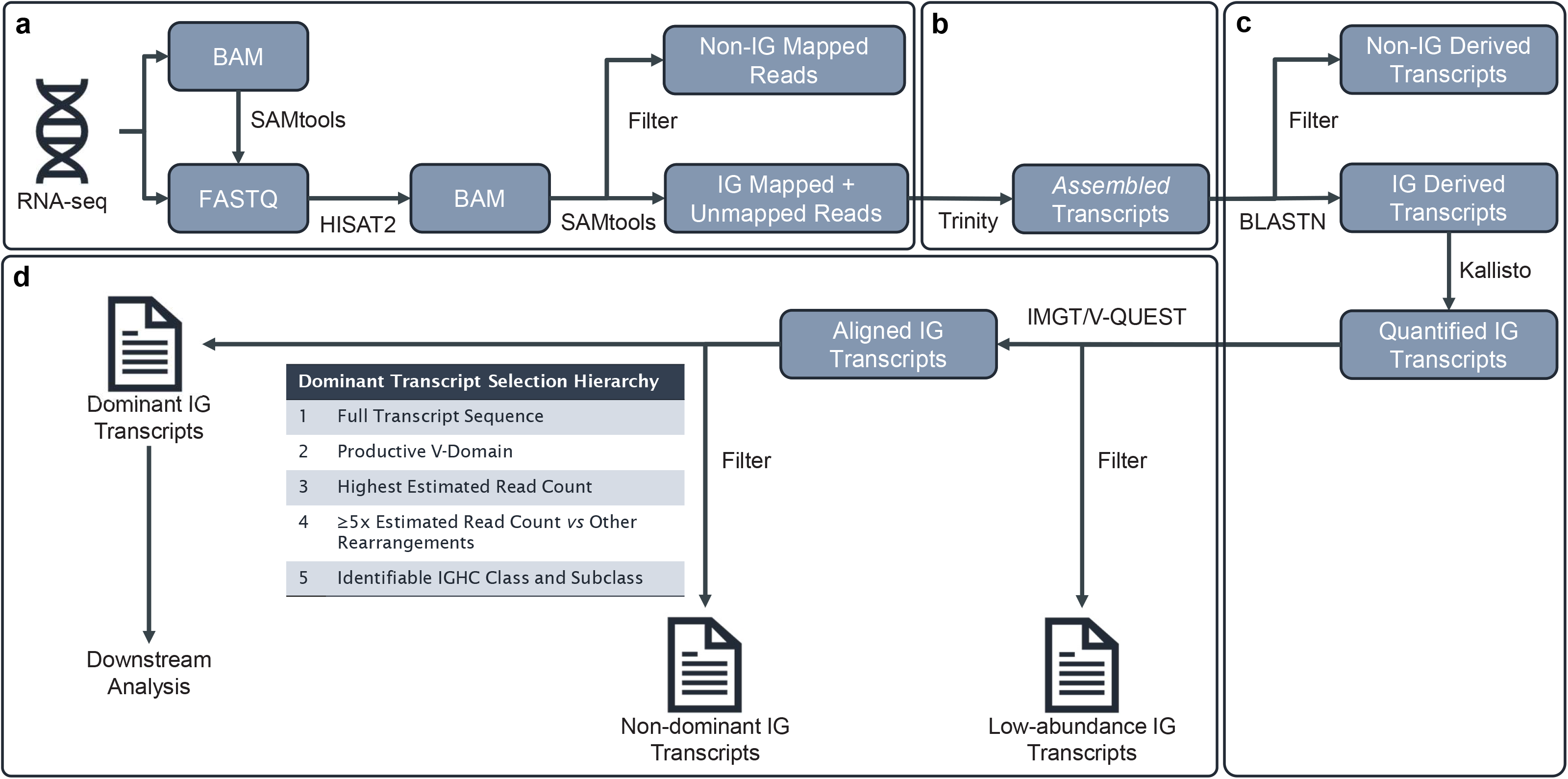
Schematic representation of the IgSeqR Pipeline. The experimental design of IgSeqR is divided into four key stages: (***a***) data pre-processing – RNA sequencing data (RNA-seq) can be supplied in either BAM or FASTQ format. The data are re-aligned to a reference transcriptome by HISAT2, producing a BAM file which is filtered to retain reads mapping to *IG* gene coordinates, and reads unable to be mapped to the reference; (***b***) *de novo* transcriptome assembly - Trinity is used to *de novo* assemble transcripts from the filtered BAM file; (***c***) *IG* transcript selection and quantification – the assembled transcripts are run through a BLAST query to identify transcripts overlapping *IG* reference sequences. The abundance of the *IG*-derived transcripts is then estimated using Kallisto pseudoalignment; (***d***) *IG* transcript annotation and interpretation – the five most abundant transcripts by TPM are then run through IMGT/V-QEUEST for *IG* alignment and annotation which is used to recover the putative tumor/dominant *IG* transcript using a 5 step hierarchical selection process.

When efficiency was assessed, IgSeqR took on average 1.18 seconds per nucleotide assembled (s/nt) to complete, compared to 8.10 s/nt and 1.44 s/nt minutes by MiXCR and TRUST4, respectively (**Table S3**).

Overall, IgSeqR obtained longer transcripts, was more efficient per nucleotide assembled, and was more accurate than MiXCR and TRUST4.

### Experimental Design

The experimental design of IgSeqR is divided into four key stages (**Figure 3**): (*a*) data preprocessing, (*b*) *de novo* transcriptome assembly, (*c*) *IG* transcript selection and quantification, and (*d*) *IG* transcript annotation and interpretation.

## Data preprocessing

IgSeqR can use RNA-seq data in either BAM or FASTQ format. We have assessed the quality of the RNA-seq data using FastQC [41], but alternative methods more familiar to the operator can be used. Alignment of the data to a reference transcriptome is performed using HISAT2 alignment tool, which employs a hierarchical indexing strategy based on Burrows-Wheeler Transform [42]. If the input file has been previously aligned, FASTQ reads must first be extracted from the alignment file (Step 1) before being supplied to HISAT2 (Step 2). It is problematic to map IG variable genes, especially D and J to a reference transcriptome using short read RNA-seq data, which results in many *IG*-derived reads being unmapped following alignment [32]. Therefore, following alignment, the resultant BAM file is filtered to extract reads which align to specific *IG* associated genomic loci in addition to any unmapped reads.

### *De Novo* Assembly

The Trinity software [43] is used for reference-free transcript reconstruction of the reads associated with *IG* sequences. Trinity follows a three-step process: Inchworm, Chrysalis, and Butterfly [43]. Inchworm builds initial contigs by assembling overlapping k-mers from the short reads. Chrysalis constructs a De Bruijn graph using the Inchworm contigs to represent connections between overlapping sequences and identifies alternative splicing events. Butterfly decomposes the De Bruijn graph into individual components representing distinct transcripts from the same gene. These components are refined and merged to generate complete transcript sequences. The filtered FASTQ files generated from the HISAT2 output are supplied to Trinity for *de novo* transcript assembly, resulting in a FASTA file containing the assembled transcripts (Step 4).

### *IG* Transcript Selection and Quantification

To remove any non-*IG* associated transcripts assembled by Trinity, the transcripts in the output FASTA file are aligned to reference *IG* databases using BLAST [44]. Reference FASTA *IG* sequences are concatenated to generate the databases (Step 5). Transcripts that align with an *IG* reference sequence are retained (Step 6) and quantified using Kallisto [45], a tool that quantifies transcript abundance from RNA-Seq data using pseudo-alignment instead of read alignment. A k-mer-based index is built (Step 7) for quantification of the filtered transcripts using the FASTQ reads (Step 8).

### *IG* Transcript Annotation and Interpretation

The most abundant transcripts are identified using the transcript quantification outputs (Step 9) and passed through the IMGT/V-QUEST sequence alignment web tool [46], benefiting from a comprehensive database of known germline *IG* alleles and polymorphisms for functional annotations (Step 10). V-QUEST identifies and annotates *IGHV-IGHD-IGHJ* and *IGKV-IGKJ* or *IGLV-IGLJ* rearrangements, detects nucleotide mutations and insertions/deletions, and functionality. The annotated transcripts are then manually reviewed to identify the tumor transcript through a hierarchical filtering process (Step 11).

### Expertise Required

To effectively implement IgSeqR, individuals must be familiar with computational biology and have basic expertise in navigating a Linux command-line environment. Users will need to be comfortable installing the necessary bioinformatics tools involved, preferably via the conda package manager. The protocol provides annotated scripts to run the pipeline, although proficiency in scripting languages, particularly BASH, and large-scale sequencing data and their data formats (**Table 3**) is beneficial. Familiarity with the principles of immunogenetics, BCR structure and function, and B-cell biology in health and disease is expected for the interpretation and curation of the results (https://www.imgt.org/IMGTeducation/). While the protocol can be performed by a skilled graduate student or postdoctoral researcher with the necessary computational expertise, collaboration with a specialized core facility for sequencing analysis may be advantageous when generating and processing primary high-throughput sequencing data.

**Table 3.** List of acronyms used in the IgSeqR protocol.

| **Acronyms** | **Name** | **Description** |
| --- | --- | --- |
| BAM | Binary Alignment/Map | The BAM format is a binary representation of sequence alignment data. It is commonly used in genomics to store the results of sequence alignment algorithms. |
| BASH | Bourne Again Shell | Unix shell and a command language interpreter. It is a default command interpreter on most GNU/Linux systems. Bash can also read and execute commands from a file, called a shell script. |
| CPU | Central Processing Unit | The CPU is the most important processor in a given computer, responsible for performing basic arithmetic, logic, controlling, and input/output (I/O) operations specified by the instructions in a program |
| CSV | Comma-Separated Values | CSV is a simple file format used to store tabular data, such as a spreadsheet or database. Each line of the file represents a row of the table, and the values are separated by commas. CSV files are widely supported by spreadsheet and database software, making them easy to import and export data. |
| FASTA | FASTA Sequence Format | The FASTA format is a text-based format for representing nucleotide or protein sequences. It consists of a single-line description followed by lines of sequence data. The format is widely used in bioinformatics for storing and exchanging sequence data. |
| FASTQ | FASTQ Sequence Format | The FASTQ format is a text-based format for storing both a biological sequence (usually nucleotide sequence) and its corresponding quality scores. It is widely used to represent raw sequencing data from high-throughput sequencing platforms. |
| TSV | Tab-Separated Values | TSV is a file format similar to CSV, but with tab characters as the field separator instead of commas. TSV files are commonly used for storing and exchanging tabular data, especially when the data may contain commas or other special characters. |
| RAM | Random Access Memory | A temporary memory bank in a computer where data which requires quick access is stored. It keeps data easily accessible so a computers processor can quickly find it without having to go into long-term storage to complete immediate processing tasks. |

### Limitations of Method

Our initial use of IgSeqR with RNA-seq data from a cohort with unknown tumor B cell percentage [20, 21] demonstrated the utility of our protocol [14]. We used selection criteria that were designed to have the maximal confidence that the sequence identified was tumor-derived (at least 5-fold higher frequency than any other functional full transcript with different CDR3 identified). The full tumor *IGHV-IGHD-IGHJ* sequences including the *IGHC* constant region isotype were defined in 339/489 (69%) samples with RNA-seq data available [14]. However, the probability of identifying the tumor sequence could be maximized by changing certain parameters, including the fold increase of the dominant to the other sequences’ frequency or the length of the transcript desired.

Although the success rate was lower compared to those with ≥10% estimated B cells, a full IG rearrangement could be identified in many samples with <10%.

The main limitation of IgSeqR is accessibility to high-quality RNA and the experimental costs of RNA-seq. However, the costs might be a limitation for large-scale cohorts, and not for well-selected samples. The poor quality of the RNA-seq data is a limitation. RNA extracted from formalin-fixed paraffin-embedded (FFPE) tumor samples, which are commonly available in diagnostic settings, is often of low quality [47], and is currently inadequate for IgSeqR.

The sequencing chemistry employed during data generation can influence the outputs of the protocol. IgSeqR protocol has been designed and tested using paired-end sequencing, which is recommended for *de novo* assembly of RNA libraries generated from a polyA library prep and allows the recovery of unmapped reads [48]. The use of sequencing assays and analytical pipelines that remove unmapped reads will severely limit IgSeqR reliability and should therefore not be used.

A benchmarking comparison of 10 DLBCL samples demonstrated notably longer runtimes when compared to our CLL cohort, with average runtimes taking 247 minutes in DLBCL *vs* 33 minutes in CLL per sample (**Table S4**). The cellular complexity and lower tumor purity (**Table S1**) of a DLBCL tissue sample may contribute to these longer runtimes compared to CLL blood samples. However, this is likely to have been compounded by the higher number of starting reads in DLBCL cases (121.2 million on average) compared to CLL (71.1 million on average), which increases the processing requirements at each stage of the protocol.

Overall, sample characteristics, sequencing chemistry, and data quality may limit the efficacy of IgSeqR. Quality control assessments should be performed, and any necessary errors should be corrected before using IgSeqR.

## Materials

### Hardware

The IgSeqR protocol is designed to be versatile, allowing compatibility with various computing resources, ranging from laptops to high-performance computing clusters, and cloud computing platforms. All analyses, including those for comparison with MixCR and TRUST4, were conducted using the Iridis5 high-performance computing cluster at the University of Southampton, utilizing 8 × 2.0 GHz CPU cores and 32 GB RAM to simulate a typical desktop workstation. Default settings were used for MixCR and TRUST4 following the RNA-seq from raw FASTQ files protocols from each tool’s documentation.

However, the protocol can be run on less powerful hardware with longer expected runtimes. Before starting the protocol, users should carefully consider the exact resources available on their machine, including CPU cores and RAM (considering the RAM utilized by the operating system), to mitigate errors.

### Software

- Operating system: Linux distribution (tested on Red Hat Enterprise v 7.9 and Ubuntu versions 16, 22 and 24 distributions)
- Conda package manager (https://conda.io) to install the IgSeqR environment. All dependencies of IgSeqR are documented in the environment file (**Supplement 1**), which eliminates the need for manual installation of individual tools and dependencies. The main software tools used in IgSeqR are listed below along with their versions as documented in the environment file:
  ∘ BLAST (v 2.13.0) [44]
  ∘ HISAT2 (v 2.2.1) [42]
  ∘ Kallisto (v 0.48.0) [45]
  ∘ Samtools (v 1.16.1) [49]
  ∘ Trinity (v 2.13.2) [43]

To create a conda environment from the command line, navigate to the directory containing the environment file and run the following command:

~~~
$ conda env create -f environment.yml
~~~

Replacing ‘`environment.yml`’ with the filepath of the environment file.

Once the environment is created, it can be activated by running the following command:

~~~
$ conda activate IgSeqR
~~~

- The protocol below provides a detailed explanation of each command required for the operation of the IgSeqR protocol. Each command can be run independently; however, the protocol is designed to be run as a complete pipeline from a Linux shell script. An example BASH script has been provided (**Supplement 2**) which will carry out all analytical steps, if a conda evironment containing the necessary software (described above) is correctly setup and the correct experimental variables have been included in the accompanying configuration file (**Supplement 3**). This can be performed by calling the following command in the directory outputs and intermediate files should be written to:

~~~
$ bash path/to/IgSeqR/Script.sh
~~~

Where ‘`path/to/IgSeqR/Script.sh`’ specifies the location of the IgSeqR script file (**Supplement 2**)

Users must read the protocol thoroughly before performing analysis using the provided scripts to facilitate error debugging and configuration for experiment-specific requirements.

### Data

In order to implement this protocol users will need:

- Paired-end RNA sequencing data in either FASTQ or BAM format
- Indexed reference transcriptome for HISAT2 alignment. The protocol was designed and tested using the HISAT2 pre-indexed GRCh38 reference which can be downloaded from the HISAT2 Repository using the command:

~~~
$ wget https://genome-idx.s3.amazonaws.com/hisat/grch38_snptran.tar.gz
~~~

Alternatively, custom indexed reference from a user provided reference transcriptome can be generated using the `hisat2-build` command, as described in the HISAT2 documentation (https://daehwankimlab.github.io/hisat2/manual/)

- Genomic coordinates associated with target regions for *de novo* transcript assembly. In this application, we have focused on *IG* heavy and light chains coordinateswhich are supplied in the Procedure section below.
- Reference sequences for IG heavy (*IGHV, IGHD, IGHJ*) and light (*IGKV, IGKJ, IGLV, IGLJ*) chain genes. The references used to develop this protocol can be found in the supplementary material (**Supplements 4-5**) However, the IMGT database is regularly updated online. Therefore the individual gene reference files should be downloaded from IMGT (**Table 4**) and merged into reference FASTA files for IG heavy and IG light chains before use of the pipeline.

**Table 4.** Links to the most recent IMGT/V-QUEST reference immunoglobulin heavy and light chain FASTA sequences.

| Chain | Gene | IMGT Link |
| --- | --- | --- |
| Heavy | IGHV | imgt.org/download/V-QUEST/IMGT_V-QUEST_reference_directory/Homo_sapiens/IG/IGHV.fasta |
|  | IGHD | imgt.org/download/V-QUEST/IMGT_V-QUEST_reference_directory/Homo_sapiens/IG/IGHD.fasta |
|  | IGHJ | imgt.org/download/V-QUEST/IMGT_V-QUEST_reference_directory/Homo_sapiens/IG/IGHJ.fasta |
| Light | IGKV | imgt.org/download/V-QUEST/IMGT_V-QUEST_reference_directory/Homo_sapiens/IG/IGKV.fasta |
|  | IGKJ | imgt.org/download/V-QUEST/IMGT_V-QUEST_reference_directory/Homo_sapiens/IG/IGKJ.fasta |
|  | IGLV | imgt.org/download/V-QUEST/IMGT_V-QUEST_reference_directory/Homo_sapiens/IG/IGLV.fasta |
|  | IGLJ | imgt.org/download/V-QUEST/IMGT_V-QUEST_reference_directory/Homo_sapiens/IG/IGLJ.fasta |

## Procedure

### Data pre-processing of newly generated sequencing data (Pre-pipeline)

**1.1** The pipeline has been optimized on fastq files from Illumina sequencing platforms (Illumina, Hayward, CA, USA). Users with newly generated sequencing data in BCL format should follow illlumina protocols for converting data into fastq format. Users with fastq files should commence the pipeline at “Step 2. *Genome Alignment*”

### Pre-processing published and existing sequencing data (∼ 5 minutes)

**1.2** FASTQ files are required for downstream steps in this pipeline, however published RNA-seq datasets often provide aligned or unaligned BAM files, in which case FASTQ records must first be extracted from these files, using the `fastq` command from Samtools.

CRITICAL STEP: The `fastq` command requires BAMs to first be sorted by name rather than the default sorting by chromosomal coordinates to ensure proper read pairing. This can be achieved by running the Samtools `sort` command.

The following example command could be used to sort, and extract FASTQ records from a paired-end BAM file ‘`sample.bam`’. This command uses 8 CPU threads for parallelization, and outputs compressed FASTQ files for read 1, read 2, and unpaired singleton reads to ‘`read1.raw.fastq.gz`’, ‘`read2.raw.fastq.gz`’, respectively:

~~~
$ samtools sort -n -@ 8 sample.bam -o sorted.bam
$ samtools fastq -@ 8 -n -c 6 sorted.bam \
-1 read1.raw.fastq.gz \
-2 read2.raw.fastq.gz \
-0 /dev/null -s /dev/null
~~~

The `-n` parameter in `sort` is used to sort BAM file by name, -@ parameter specifies the number of CPU threads to be used for parallelization of tasks. The `- n` option in `fastq` is used to leave the read names as they are provided. The `-c` option sets the compression level of the output files. ‘`sample.bam’` specifies the path to the input BAM file. `-1` and `-2` specify the desired paths for the compressed FASTQ output files for read 1 and read 2, respectively. `-0 /dev/null` and `-s /dev/null` discard any discarding singletons, supplementary and secondary reads.

CRITICAL STEP: It is important to perform quality control (QC) to ensure that the data is of sufficient quality for downstream analysis. A widely used QC tool is FastQC, which produces a detailed report of several quality metrics including per base sequence quality, per sequence quality scores, per base sequence content, per sequence GC content, and sequence length distribution, among others detailed at in the FastQC documentation [41]. If any issues are identified, corrective measures should be taken as per local procedures or general best practice [50]. If the tumor purity is unknown, it is advisable to estimate this through identification of the B cell proportion using a computational cellular deconvolution tool such as Cibersort [31].

### Genome Alignment (∼ 10 minutes)

**2.** FASTQ reads are aligned to a reference genome using HISAT2 which produces SAM output file which is processed by Samtools. These commands can be run as a pipeline to save computational resources. The HISAT2 SAM can be passed to Samtools `view` for conversion to BAM format which is then sorted using the Samtools `sort` command. Upon completion of Samtools `sort,` Samtools `index` is run to create an accompanying index file for the BAM.

The following example command can be used to align FASTQ input files ‘`read1.raw.fastq.gz`’ and ‘`read2.raw.fastq.gz`’ to the GRCh38 reference transcriptome using 8 CPU threads. The resulting HISAT2 aligned BAM file is output as ‘`hisat_output.bam`’ and its corresponding index as ‘`hisat_output.bam.bai`’:

~~~
$ hisat2 -p 8 --phred33 -x grch38_snp_tran \
-1 read1.raw.fastq.gz -2 read1.raw.fastq.gz | \
samtools view -@8 -bS -0 - - | \
samtools sort -@8 - -o hisat_output.bam &&
samtools index -@8 hisat_output.bam -o hisat_output.bam.bai
~~~

The `-p` or -@ parameters specifies the number of CPU threads to be used for parallelization, while `--phred33` specifies the encoding format of the quality scores. The `-x` parameter specifies the path and basename of the indexed reference transcriptome files. The input FASTQ file paths are specified by `-1` and `-2` for read 1 and read 2, respectively. The SAM is converted to bam using `-bS` with `-0` specifying no additional filtering or format conversions, and `-` signifying the standard input from the previous command. The sorted output is written to a file path specified by `-o hisat_output.bam` from which an index file is created and written to the file path specified by `–o hisat_output.bam.bai`.

### Read selection (<5 minutes)

**3.** Samtools is used to remove all reads except those that map to the *IG*-associated loci and those that are unmapped from the HISAT2 aligned BAM file, ensuring that highly variable *IG* regions that are difficult to map are retained.

TROUBLESHOOTING: If working from existing data, unmapped reads may have been removed and the alignment files may not contain sufficient IG reads to produce quality results.

The `view` command is used to extract the *IG*-associated loci and unmapped reads independently, before joining using the `merge` command.

The following example command can be used to filter the HISAT2 aligned BAM file ‘`hisat2_output.bam`’, retaining reads mapping to the *IGH, IGK*, and *IGL* loci and unmapped reads, using 8 threads for parallelization. The resulting filtered bam BAM file is output as ‘`IG_filtered.bam`’. Process substitution can be applied when using a supported Unix shell to avoid the generation of temporary files:

~~~
$ samtools merge -f IGH_filtered.bam \
<(samtools view -@ 8 -b -f 4 hisat2_output.bam) \
<(samtools view -@ 8 -b hisat2_output.bam 14:105550000-
106900000 2:87000000-92000000 22:20500000-24500000)
~~~

Where -@ specifies the number of CPU threads to be utilized for parallelization, `-b` specifies the output format as BAM, `-f 4` returns sequences which have the unmapped Samtools flag, `hisat2_output.bam` is the full input HISAT2 aligned BAM file. The *IG* coordinates `14:105550000-106900000, 2:87000000-92000000` and `22:20500000-24500000` for *IGH, IGK* and *IGL*, respectively, are specified in the format `chr:start-end` where `chr` is the chromosome number, `start` is the numerical position of the first nucleotide in the loci and `end` is the numerical position of the last nucleotide.

TROUBLESHOOTING: The format of the *IG* coordinates will depend on the reference transcriptome used to generate the aligned BAM file. The HISAT2 indexed GRCh38 reference uses numerical values for chromosome (e.g., 14). However, other references may also include a ‘chr’ prefix (e.g., chr14). Additionally, if a different reference transcriptome build is used (e.g., GRCh37) the coordinates should be converted accordingly.

### De Novo Transcript Assembly (∼ 15 minutes)

**4.** Trinity accepts FASTQ input files which must be extracted from the ‘`IGH_filtered.bam`’ BAM file using the Samtools `fastq` command (as described in Step 1.2).

The following example command can be used to perform Trinity *de novo* assembly with the input filtered FASTQ files ‘`IG_filtered_read1.fastq`’ and ‘`IG_filtered_read2.fastq`’, using 8 threads for parallelization and 32Gb RAM. The resulting transcriptome FASTA file is output as ‘`trinity_transcripts.fasta`’:

~~~
$ Trinity –CPU 8 –max_memory 32G –seqType fq \
--left IG_filtered_read1.fastq \
--right IG_filtered_read2.fastq \
--output trinity_transcripts \
--no_normalize_reads \
--min_contig_length 500 \
--full_cleanup
~~~

Where `--CPU` specifies the number of CPU threads to be utilized for parallelization, `--max_memory` specifies the maximum memory to be utilized, `-- seqType fq` specifies that the input files are in FASTQ format, `--left` and `– right` are the filtered input FASTQ files for read 1 and read 2, respectively, and `–output <output>` is the basename of the output files.

CRITICAL STEP: Read normalization aims to reduce bias in assembly by down sampling highly expressed reads. Input data will be enriched for *IG* transcripts. This can lead to a reduction of reads for low-abundance transcripts, which can lead to incomplete assembly or loss of rare transcripts and should be disabled using `– no_normalize_reads`.

CRITICAL STEP: Short contigs may represent partial or fragmented *IG* transcripts, which can affect downstream analysis and interpretation. Using a minimum contig length of 500 with `–min_contig_length`, most assembled transcripts will contain the full *IGV-(IGD)-IGJ* recombination.

### IG Transcript Selection (< 5 mins)

The protocol permits the detection and quantification of *IG* heavy and/or light chains. The steps below provide examples of *IG* heavy chain transcript extraction, but can be adapted to extract the *IG* light chain transcript.

**5.** To extract putative *IG* sequences from the Trinity assembly, the transcriptome FASTA file containing the assembled contigs are searched against a reference sequence using BLAST.

Individual BLAST databases should be generated using the reference sequences for *IG* heavy (*IGHV,IGHD,IGHJ*) (**Supplement 4**) and light (*IGKV,IGKJ,IGLV,IGLJ*) (**Supplement 5)** chains as required using the `makeblastdb` command.

The following example command describes how to generate a BLAST database from the IG heavy reference FASTA sequences ‘`IGH_reference.fasta`’:

~~~
$ makeblastdb -in IGH_reference.fasta -parse_seqids -dbtype nucl
~~~

Where `-in` specifies the input FASTA file containing reference sequences, `-parse_seqids` allows the FASTA headers to be parsed along with their sequence, and `-dbtype nucl` specifies the sequence content to be nucleotides.

TROUBLESHOOTING: Ensure that the sequence headers in the FASTA file do not contain the pipe (“|”) character as it is a reserved character for the ID parser, which can cause an error.

**6.** The assembled transcripts are compared against the reference database(s) generated in step 5 using the `BLASTN` command. This produces a tabular output that can be passed to the `cut` and `uniq` commands to obtain a unique list of *IG* transcript IDs, which are used by `samtools faidx` to extract the corresponding sequences from the assembled transcripts FASTA file.

The following example command can be used to select *IG* transcripts covering reference *IG* heavy FASTA sequences in the ‘`IGH_reference.fasta`’ file from the assembled transcripts ‘`trinity_transcripts.fasta`’, to produce the filtered FASTA file ‘`IGH_transcripts.fasta`’:

~~~
$ blastn -db IGH_reference.fasta \
-query trinity_transcripts.fasta -outfmt 6 | \
cut -f1 | uniq | xargs -n 1 samtools faidx
trinity_transcripts.fasta > IGH_transcripts.fasta
~~~

`Where, -db` specifies the path to the FASTA file used to generate the reference database for either *IG* heavy or light sequences, `-query` specifies the path to the FASTA file containing the Trinity assembled transcripts, `-outfmt 6` sets the output format to be tabular, `cut -f1` selects the transcript ID (first) column in the tabular `BLASTN` output, `uniq` removes duplicate transcript IDs, `xargs -n 1` reads the IDs from output of the `uniq` (one ID per line) and passes them to `samtools faidx` as separate arguments.

### Transcript Quantification (< 5 minutes)

**7.** Abundance of selected transcripts is quantified using the Kallisto pseudoalignment tool which first requires a Kallisto index to be built from the input FASTA file using the `index` command.

The following example command can be used to generate a Kallisto index file ‘`kallisto.index`’ for the *IG* heavy chain filtered transcript FASTA sequence file ‘`IGH_transcripts.fasta`’:

~~~
kallisto index -i kallisto.index IGH_transcripts.fasta
~~~

Where `-i` specifies the filename of the Kallisto index to be constructed and ‘`IGH_transcripts.fasta`’ is the path to the filtered *IG* transcripts FASTA sequences.

**8.** The generated index is used in the `quant` command, along with FASTQ files used to assemble the transcripts to quantify the abundance of the *IG* filtered transcripts.

The following example command can be used to quantify the abundance of transcripts in the *IG* filtered transcript FASTQ files (generated in step 4) ‘`IG_filtered_read1.fastq`’ and ‘`IG_filtered_read1.fastq`’, using 8 threads:

~~~
kallisto quant -i kallisto.index -t 8 \
IG_filtered_read1.fastq IG_filtered_read2.fastq
~~~

Where `-i` specifies the filename of the Kallisto index, `-t` specifies the number of CPU threads to be utilized for parallelization, and ‘`IG_filtered_read1.fastq`’ and ‘`IG_filtered_read2.fastq`’ are the *IG* filtered FASTQ files for read 1 and read 2, respectively.

**9.** The five most abundant transcripts IDs are identified based on their transcript per million (TPM) value by passing the Kallisto output through the `tail`, `sort`, `head` and `cut` commands, and their corresponding FASTA sequences are extracted using `samtools faidx command`.

The following example command can be used to identify the five most abundant transcript IDs from the Kallisto output ‘`abundance.tsv`’, extract their corresponding transcript sequences from ‘`IGH_transcripts.fasta`’ and write to an output FASTA file called ‘`IGH_TPM_filtered.fasta`’:

~~~
$ tail -n +2 abundance.tsv | \
sort -t $’\t’ -k5,5nr | head -5 | cut -f1 | \
xargs -n 1 samtools faidx IGH_transcripts.fasta >
IGH_TPM_filtered.fasta
~~~

Where `-n +2` selects all rows except the first (header) from the Kallisto quantification output, `-t $’\t’` specifies the delimiter of the input as tab, `- k5,5nr sorts` the remaining lines by the fifth column (TPM) in reverse numerical order, `head -5` outputs the first 5 lines of the sorted file and `cut -f1` extracts the first column (IDs) from the output. The IDs are read (one ID per line) using `xargs -n 1` which then passes them to `samtools faidx` as separate arguments.

TROUBLESHOOTING: The number of most abundant transcripts to take forward has been suggested as 5. This has been found to strike a good balance between analytical efficiency and identification of the dominant tumor transcript. In instaces where no full-length, productive transcripts are not obtained within the top 5 transcripts, users may wish to increase the number of transcripts to take forward for analysis.

### Dominant IG Transcript selection (∼ 15 minutes)

**10.** The top 5 most abundant transcripts identified in step 9 will be submitted to the IMGT/V-QUEST tool (https://imgt.org/IMGT_vquest/input) for sequence analysis and annotation. In the sequence submission section of the IMGT/V-QUEST tool, the top 5 transcript sequences should be provided either by copy and pasting the sequences from the FASTA file or by directly uploading the FASTA file. The parameters ‘Species’ and ‘Receptor type or locus’ should be set to ‘Homo sapiens (human)’ and ‘IG’, respectively. Finally, the output format should be set to ‘C.Excel file’. The IMGT/V-QUEST tool will annotate and analyze the submitted sequences for their corresponding *IGV, IGHD* (for the heavy chain only) and IGJ genes, their junction at the *CDR3* region, and other related features.

**11.** The outputs of the Kallisto quantification and IMGT/V-QUEST results transcript are used to identify the dominant/consensus (Tumor) *IG* transcript present within the RNA-seq dataset. This process may require manual interpretation but follows the following hierarchical filtering criteria:

i. Presence of a full transcript sequence (from codon 1 in FR1 to codon 129 in FR4 included), identified by IMGT/VQUEST.
ii. Presence of ‘productive’ V-domain functionality call by IMGT/V-QUEST
iii. The highest estimated read count (est. count) determined by Kallisto.
iv. The est. count is greater than 5-fold higher than any of the other 4 transcripts selected if different. A reduction of the fold amount difference will increase the probability to identify a “dominant” sequence in cases with low tumor infiltration.
v. The ability to determine the *IG* constant region class and subclass.

### Timing

Benchmarking was conducted using the computational hardware described in the materials section. The dominant *IG* heavy and light chain transcripts were extracted from FASTQ files generated from high-purity CLL samples with an average starting read count of 71.1 million following initial HISAT2 alignment. In similar conditions, the full pipeline can be expected to take less than 1 hour per sample. Specific timings can be found in the procedure section headers for each stage of the analytical pipeline. The duration of each stage may vary depending on the input file type (BAM files require additional pre-processing), hardware used to run the pipeline, heterogeneity of B-cell populations, and number of starting sequencing reads generated from the samples.

### Anticipated results

Upon successful completion of the IgSeqR protocol, users will have generated the following output files for *IG* heavy and/or light chain transcripts:

- The five most abundant assembled *IG* transcripts in FASTA format
- Table of quantifications for these *IG* transcripts in tsv format
- Annotations for the top five *IG* transcripts generated by IMGT/V-QUEST.

For further insights, users can refer to our previously published [14], which includes results and examples of downstream analysis.

## Future Applications

Future work is planned to develop further the existing protocol and evaluate its efficacy for deriving smaller, less dominant, clonal populations to widen the application of the protocol. When the tumor *IG* sequence is already known, we will apply this approach for the determination of the minimal residual disease in repeat samples following anti-cancer therapy.

We will also investigate the protocol’s potential use with RNA-seq data generated from FFPE material. However, there are intrinsic limitations of RNA-seq data quality from FFPE, and areas where optimization or adaptation may be necessary will need to be identified.

Future work will also focus on the annotation refinement of the *IGC* region. This work will facilitate and accompany the development of the protocol into a comprehensive bioinformatics tool for immunobiologists.

The protocol will also be investigated for its use in any other genomic regions that are challenging to map to a reference genome, including the T-cell Receptoror specific fusion or deregulating gene rearrangements that are not represented in the reference transcriptome [51].

## Supplementary information

- Supplementary Tables.xlsx
- Supplement 1. IgSeqR Environment.yml
- Supplement 2. IgSeqR BASH Script.sh
- Supplement 3. IgSeqR Configuration File
- Supplement 4. IGH References.fasta
- Supplement 5. IGKL References.fasta

## Author contributions statements

D.B. and B.J.S. designed IgSeqR bioinformatic pipeline, analyzed and interpreted data, and wrote the manuscript. D.T. and G.C analyzed, interpreted data, and contributed to the immunoglobulin gene analysis pipeline validation. B.S., A.O. and J.B contributed to the analysis and interpretation of the data. F.F. designed the study, supervised research, interpreted data and wrote the manuscript. All authors reviewed and approved the manuscript.

## Acknowledgments

The authors are grateful to the Faculty of Medicine Tissue Bank (Cancer Sciences, University of Southampton) for the processing and storage of the primary lymphoma specimens. This work was supported by Cancer Research UK (ECRIN-M3 accelerator award C42023/A29370, and BTERP project C36811/A29101). D.T. was funded by the Eyles Cancer Immunology PhD scholarship. G.C. was funded by the Eyles Cancer Immunology Fellowship and the Southampton Cancer Immunology Centre Pump-priming award 2021). Genetic data for IgSeqR protocol were obtained via the National Cancer Institute Genomic Data Commons for Genotypes and Phenotypes (accession phs001444.v1.p1). The authors acknowledge the use of the IRIDIS High Performance Computing Facility, and associated support services at the University of Southampton, in the completion of this work.

## Competing interests

The authors declare no potential conflicts of interest.

## Notes

### Competing Interest Statement

The authors have declared no competing interest.

## References

1. Lam, K.P., R. Kuhn, and K. Rajewsky, In vivo ablation of surface immunoglobulin on mature B cells by inducible gene targeting results in rapid cell death. Cell, 1997. 90(6): p. 1073–83.

2. Stevenson, F.K., et al., The occurrence and significance of V gene mutations in B cell-derived human malignancy. Adv Cancer Res, 2001. 83: p. 81–116.

3. Victora, G.D. and M.C. Nussenzweig, Germinal Centers. Annual Review of Immunology, 2022. 40(1): p. 413–442.

4. Forconi, F., S.A. Lanham, and G. Chiodin, Biological and Clinical Insight from Analysis of the Tumor B-Cell Receptor Structure and Function in Chronic Lymphocytic Leukemia. Cancers (Basel), 2022. 14(3).

5. Stevenson, F.K., F. Forconi, and T.J. Kipps, Exploring the pathways to chronic lymphocytic leukemia. Blood, 2021. 138(10): p. 827–835.

6. Efremov, D.G., S. Turkalj, and L. Laurenti, Mechanisms of B Cell Receptor Activation and Responses to B Cell Receptor Inhibitors in B Cell Malignancies. Cancers, 2020. 12(6): p. 1396.

7. Forconi, F., et al., The normal IGHV1-69-derived B-cell repertoire contains stereotypic patterns characteristic of unmutated CLL. Blood, 2010. 115(1): p. 71–7.

8. Seifert, M., et al., Cellular origin and pathophysiology of chronic lymphocytic leukemia. J Exp Med, 2012. 209(12): p. 2183–98.

9. Damle, R.N., et al., Ig V gene mutation status and CD38 expression as novel prognostic indicators in chronic lymphocytic leukemia. Blood, 1999. 94(6): p. 1840–7.

10. Hamblin, T.J., et al., Unmutated Ig V(H) genes are associated with a more aggressive form of chronic lymphocytic leukemia. Blood, 1999. 94(6): p. 1848–54.

11. Niemann, C.U., et al., Fixed-duration ibrutinib–venetoclax versus chlorambucil– obinutuzumab in previously untreated chronic lymphocytic leukaemia (GLOW): 4-year follow-up from a multicentre, open-label, randomised, phase 3 trial. The Lancet Oncology, 2023.

12. Stevenson, F.K. and F. Forconi, The essential microenvironmental role of oligomannoses inserted into the antigen-binding sites of lymphoma cells. Blood, 2023.

13. Zhu, D., et al., Acquisition of potential N-glycosylation sites in the immunoglobulin variable region by somatic mutation is a distinctive feature of follicular lymphoma. Blood, 2002. 99(7): p. 2562–2568.

14. Chiodin, G., et al., Insertion of atypical glycans into the tumor antigen-binding site identifies DLBCLs with distinct origin and behavior. Blood, 2021. 138(17): p. 1570–1582.

15. Coelho, V., et al., Glycosylation of surface Ig creates a functional bridge between human follicular lymphoma and microenvironmental lectins. Proc Natl Acad Sci U S A, 2010. 107(43): p. 18587–92.

16. Linley, A., et al., Lectin binding to surface Ig variable regions provides a universal persistent activating signal for follicular lymphoma cells. Blood, 2015. 126(16): p. 1902–10.

17. Odabashian, M., et al., IGHV sequencing reveals acquired N-glycosylation sites as a clonal and stable event during follicular lymphoma evolution. Blood, 2020. 135(11): p. 834–844.

18. Sutton, L.A., et al., Immunoglobulin genes in chronic lymphocytic leukemia: key to understanding the disease and improving risk stratification. Haematologica, 2017. 102(6): p. 968–971.

19. Wang, Z., M. Gerstein, and M. Snyder, RNA-Seq: a revolutionary tool for transcriptomics. Nat Rev Genet, 2009. 10(1): p. 57–63.

20. Schmitz, R., et al., Genetics and pathogenesis of diffuse large B-cell lymphoma. New England Journal of Medicine, 2018. 378(15): p. 1396–1407.

21. Wright, G.W., et al., A Probabilistic Classification Tool for Genetic Subtypes of Diffuse Large B Cell Lymphoma with Therapeutic Implications. Cancer Cell, 2020. 37(4): p. 551-568.e14.

22. Bryant, D., et al., Network analysis reveals a major role for 14q32 cluster miRNAs in determining transcriptional differences between IGHV-mutated and unmutated CLL. Leukemia, 2023. 37(7): p. 1454–1463.

23. Teo, Q.W., et al., Stringent and complex sequence constraints of an IGHV1-69 broadly neutralizing antibody to influenza HA stem. Cell Reports, 2023. 42(11): p. 113410.

24. Shiroishi, M., Structural Basis of a Conventional Recognition Mode of IGHV1-69 Rheumatoid Factors, in Protein Reviews : Volume 21, M.Z. Atassi, Editor. 2021, Springer International Publishing: Cham. p. 171–182.

25. Forconi, F., et al., Insight into the potential for DNA idiotypic fusion vaccines designed for patients by analysing xenogeneic anti-idiotypic antibody responses. Immunology, 2002. 107(1): p. 39–45.

26. Stevenson, G.T., E.V. Elliott, and F.K. Stevenson, Idiotypic determinants on the surface immunoglobulin of neoplastic lymphocytes: a therapeutic target. Fed Proc, 1977. 36(9): p. 2268–71.

27. Hawkins, R.E., et al., Idiotypic vaccination against human B-cell lymphoma. Rescue of variable region gene sequences from biopsy material for assembly as single-chain Fv personal vaccines. Blood, 1994. 83(11): p. 3279–88.

28. McCann, K., et al., Idiotype gene rescue in follicular lymphoma. Methods Mol Med, 2005. 115: p. 145–71.

29. Ottensmeier, C.H. and F.K. Stevenson, Isotype switch variants reveal clonally related subpopulations in diffuse large B-cell lymphoma. Blood, 2000. 96(7): p. 2550–6.

30. Ottensmeier, C.H., et al., Analysis of VH genes in follicular and diffuse lymphoma shows ongoing somatic mutation and multiple isotype transcripts in early disease with changes during disease progression. Blood, 1998. 91(11): p. 4292–9.

31. Newman, A.M., et al., Robust enumeration of cell subsets from tissue expression profiles. Nature Methods, 2015. 12(5): p. 453–457.

32. Blachly, J.S., et al., Immunoglobulin transcript sequence and somatic hypermutation computation from unselected RNA-seq reads in chronic lymphocytic leukemia. Proc Natl Acad Sci U S A, 2015. 112(14): p. 4322–7.

33. Bolotin, D.A., et al., MiXCR: software for comprehensive adaptive immunity profiling. Nat Methods, 2015. 12(5): p. 380–1.

34. Canzar, S., et al., BASIC: BCR assembly from single cells. Bioinformatics, 2017. 33(3): p. 425–427.

35. Kuchenbecker, L., et al., IMSEQ--a fast and error aware approach to immunogenetic sequence analysis. Bioinformatics, 2015. 31(18): p. 2963–71.

36. Mandric, I., et al., Profiling immunoglobulin repertoires across multiple human tissues using RNA sequencing. Nat Commun, 2020. 11(1): p. 3126.

37. Mose, L.E., et al., Assembly-based inference of B-cell receptor repertoires from short read RNA sequencing data with V’DJer. Bioinformatics, 2016. 32(24): p. 3729–3734.

38. Rizzetto, S., et al., B-cell receptor reconstruction from single-cell RNA-seq with VDJPuzzle. Bioinformatics, 2018. 34(16): p. 2846–2847.

39. Song, L., et al., TRUST4: immune repertoire reconstruction from bulk and single-cell RNA-seq data. Nat Methods, 2021. 18(6): p. 627–630.

40. Upadhyay, A.A., et al., BALDR: a computational pipeline for paired heavy and light chain immunoglobulin reconstruction in single-cell RNA-seq data. Genome Med, 2018. 10(1): p. 20.

41. Andrews, S., FastQC: A Quality Control Tool for High Throughput Sequence Data. Babraham Bioinformatics, 2010.

42. Kim, D., et al., Graph-based genome alignment and genotyping with HISAT2 and HISAT-genotype. Nature Biotechnology, 2019. 37(8): p. 907–915.

43. Grabherr, M.G., et al., Full-length transcriptome assembly from RNA-Seq data without a reference genome. Nat Biotechnol, 2011. 29(7): p. 644–52.

44. Altschul, S.F., et al., Basic local alignment search tool. J Mol Biol, 1990. 215(3): p. 403–10.

45. Bray, N.L., et al., Near-optimal probabilistic RNA-seq quantification. Nat Biotechnol, 2016. 34(5): p. 525–7.

46. Brochet, X., M.P. Lefranc, and V. Giudicelli, IMGT/V-QUEST: the highly customized and integrated system for IG and TR standardized V-J and V-D-J sequence analysis. Nucleic Acids Res, 2008. 36(Web Server issue): p. W503–8.

47. Cazzato, G., et al., Formalin-Fixed and Paraffin-Embedded Samples for Next Generation Sequencing: Problems and Solutions. Genes (Basel), 2021. 12(10).

48. Haas, B.J., et al., De novo transcript sequence reconstruction from RNA-seq using the Trinity platform for reference generation and analysis. Nat Protoc, 2013. 8(8): p. 1494–512.

49. Li, H., et al., The Sequence Alignment/Map format and SAMtools. Bioinformatics, 2009. 25(16): p. 2078–9.

50. Hesketh, A.R., RNA Sequencing Best Practices: Experimental Protocol and Data Analysis, in Yeast Systems Biology: Methods and Protocols, S.G. Oliver and J.I. Castrillo, Editors. 2019, Springer New York: New York, NY. p. 113–129.

51. Haas, B.J., et al., Accuracy assessment of fusion transcript detection via read-mapping and de novo fusion transcript assembly-based methods. Genome Biology, 2019. 20(1): p. 213.

