## Supplements 1 to 5 for "IgSeqR: a protocol for the identification, assembly, and characterization of full-length tumor Immunoglobulin transcripts from unselected RNA sequencing data": Supplement 1. IgSeqR Environment.yml

```
name: IgSeqR
channels:
  - conda-forge
  - bioconda
  - defaults
dependencies:
  - blast=2.13.0=hf3cf87c_0
  - hisat2=2.2.1=hdbdd923_6
  - kallisto=0.48.0=h15996b6_2
  - samtools=1.16.1=h00cdaf9_2
  - trinity=2.13.2=hff880f7_4
```
