## Supplements 1 to 5 for "IgSeqR: a protocol for the identification, assembly, and characterization of full-length tumor Immunoglobulin transcripts from unselected RNA sequencing data": Supplement 2. IgSeqR BASH Script.sh

```
#!/bin/bash

################################### Activate Conda Environment ####################################
source activate IgSeqR
condaEnv=`which env`

################################## Load Configuration Variables ##################################
# Load the variables from the config file
source config
home_dir=$(pwd)

######################################## Check Input Files #######################################

# Check if both BAM and FASTQ files are provided
if [[ -n $bam && (-n $read1 || -n $read2) ]]; then
    echo "Error: Both BAM and FASTQ files are provided. Please provide only one type."
    exit 1
fi

# Check if only one of the FASTQ files is provided
if [[ (-n $read1 && -z $read2) || (-z $read1 && -n $read2) ]]; then
    echo "Error: Both read1 and read2 FASTQ files need to be provided."
    exit 1
fi

# If BAM file is provided, check if it exists
if [[ -n $bam ]]; then
    if [[ -e $bam ]]; then
        from_bam="TRUE"
        echo 
        echo ===============================================================================
        echo IgSeqR - EXTRACT TRANSCRIPTS FROM BAM
        echo ===============================================================================
        echo
        echo ================================= Config ======================================
        echo BAM:                $bam
        echo Sample ID:          $sample
        echo Hisat Reference:     $hisat_ref
        echo IMGT Reference:      $imgt_ref
        echo Cores:              $cores
        echo Conda Environment:  $condaEnv
        echo =============================================================================== 

        #################################### Convert Bam into FASTQ #################################### 
        ## Convert BAM reads back into FASTQ for HISAT2 alignment
        echo
        date
        echo =========================== Converting BAM to FASTQ ===========================

        mkdir ./FASTQ

        echo Sorting BAM by name ... 
        samtools sort -n -@$cores $bam -o ./FASTQ/"$sample"_sorted.bam

        echo Converting BAM to FASTQ ...
        samtools fastq -@ $cores -n -c 6 ./FASTQ/"$sample"_sorted.bam \
        -1 ./FASTQ/"$sample"_read1.fastq.gz \
        -2 ./FASTQ/"$sample"_read2.fastq.gz \
        -0 /dev/null -s /dev/null

        fastq1="./FASTQ/"$sample"_read1.fastq.gz"
        fastq2="./FASTQ/"$sample"_read2.fastq.gz"     
    else
        echo "Error: BAM file $bam does not exist."
        exit 1
    fi
fi

# If FASTQ files are provided, check if they exist
if [[ -n $read1 && -n $read2 ]]; then
    if [[ -e $read1 && -e $read2 ]]; then
        echo 
        echo ===============================================================================
        echo IgSeqR - EXTRACT TRANSCRIPTS FROM FASTQ
        echo ===============================================================================
        echo
        echo ================================= Config ======================================
        echo Read1:              $read1
        echo Read2:              $read2
        echo Sample ID:          $sample
        echo Hisat Reference:     $hisat_ref
        echo IMGT Reference:      $imgt_ref
        echo Cores:              $cores
        echo Conda Environment:  $condaEnv
        echo ===============================================================================

        fastq1=$read1
        fastq2=$read2
    else
        echo "Error: One or both of the FASTQ files $read1 and/or $read2 do not exist."
        exit 1
    fi
fi

######################################## HISAT2 Alignment ########################################
## Map to genome using hisat2, sort and index bam

mkdir ./hisat2

echo
date
echo ========================= Performing HISAT2 Alignment =========================

hisat2 -p $cores --phred33 -t -x $hisat_ref -1 $fastq1 -2 $fastq2 | \
  samtools view -@$cores -bS -o - - | \
  samtools sort -@$cores - -o ./hisat2/hisat2_output.bam &&
samtools index -@$cores ./hisat2/hisat2_output.bam -o ./hisat2/hisat2_output.bam.bai

echo
echo HISAT2 Flagstat:
samtools flagstat ./hisat2/hisat2_output.bam

############################# Clean FASTQ Files if created  #####################################
if [[ -e "./FASTQ" ]]; then
  rm -r ./FASTQ
fi

######################################### Read Selection ######################################### 
## Select unmapped reads and any reads mapping to IG loci to take forward to trinity
## Remove anything not required to assemble IG transcripts
## The fewer reads, the better as trinity is very slow

## Coordinates to keep with buffer:
##  - IGH: "14:100000000-110000000"
##  - IGK: "2:87000000-92000000"
##  - IGL: "22:20500000-24500000"

## Coordinates to keep without buffer:
##  - IGH: "14:105550000-106900000" (IGHA2 to IGHVIII-82)
##  - IGK: "2:88697000-92240000" (IGKC to IGKV1OR2 pseudogene)
##  - IGL: "22:22005000-23590000" (IGLV4-69 to IGLL1)

echo
date
echo ============================== IG Read Filtering ==============================

IGH="14:105550000-106900000"
IGK="2:88697000-92240000"
IGL="22:22005000-23590000"

samtools merge -f ./hisat2/keep_reads.bam \
<(samtools view -@ $cores -b -f 4 ./hisat2/hisat2_output.bam) \
<(samtools view -@ $cores -b ./hisat2/hisat2_output.bam $IGH $IGK $IGL)

samtools sort -n -@$cores ./hisat2/keep_reads.bam -o ./hisat2/keep_reads.bam

#################################### Convert Back into FASTQ ################################### 
## Convert filtered reads back into FASTQ for running through Trinity

echo
date
echo =========================== Converting BAM to FASTQ ===========================

mkdir ./IG_READS

samtools fastq -@ $cores -n -c 6 ./hisat2/keep_reads.bam \
-1 ./IG_READS/"$sample"_read1.fastq.gz \
-2 ./IG_READS/"$sample"_read2.fastq.gz \
-0 /dev/null -s /dev/null

###################################### Clean HISAT2 Files #####################################
rm -r hisat2

########################### Trinity genome naive trascript assembly ###########################
## here we assemble the kept reads into transcripts. 
## Option choice is important, please note the following:
##   - no digital normalisation
##   - no clipping
##   - min contig 500bp
##   - RF mode
##   - non genome mode

echo
date
echo ======================= De Novo Assembly of Transcripts =======================

mkdir ./Trinity

Trinity \
 --seqType fq \
 --max_memory $mem \
 --min_contig_length 500 \
 --full_cleanup \
 --no_normalize_reads \
 --output ./Trinity/"$sample"_Trinity \
 --left ./IG_READS/"$sample"_read1.fastq.gz \
 --right ./IG_READS/"$sample"_read2.fastq.gz \
 --CPU $cores

################################### Generate IG transcripts ##################################
## BLAST assembeled transcripts to the IMGT reference FASTA file
## Collect unique transcript IDs associated with IMGT reference

echo
date
echo ========================== IG Transcript extraction ===========================

mkdir ./igseqr/

for gene in IGH IGKL
do

echo Extracting $gene transcripts:

blastn -db $imgt_ref/$gene.fasta -query ./Trinity/"$sample"_Trinity.Trinity.fasta -outfmt 6 | \
  cut -f1 | \
  uniq | \
  xargs -n 1 samtools faidx ./Trinity/"$sample"_Trinity.Trinity.fasta > ./igseqr/"$gene"_transcripts.fasta

count=$(grep ">" ./igseqr/"$gene"_transcripts.fasta | wc -l)

echo "  - $count Transcripts Extracted"
echo

done

###################################### Clean Trinity Files #####################################
rm -r Trinity

echo
echo ==============================================================================
echo "IgSeqR - QUANTIFY, ANNOTATE AND FILTER TRANSCRIPTS"
echo ==============================================================================

################################# Quantify IGH transcripts #################################
## Quantify extracted IG associated transcripts using Kallisto pseudoalignment

echo
date
echo =========================== Kallisto Quantification ===========================

for gene in IGH IGKL
do
echo "Running Kallisto for $gene Transcripts..."

mkdir -p ./igseqr/kallisto/"$gene"

## Build kallisto index for transcripts
kallisto index -i ./igseqr/kallisto/"$gene"/"$gene"_transcripts.index ./igseqr/"$gene"_transcripts.fasta

## Quantify number of reads mapping to transcripts
kallisto quant \
  -i ./igseqr/kallisto/"$gene"/"$gene"_transcripts.index \
  -o ./igseqr/kallisto/"$gene" \
  -t $cores \
  ./IG_READS/"$sample"_read1.fastq.gz ./IG_READS/"$sample"_read2.fastq.gz

done

###################################### Clean IG Read Files #####################################
rm -r IG_READS

source config
  
for gene in IGH IGKL
  do
    if [ -d "igseqr/kallisto/$gene" ]; then
      cd igseqr/
      tail -n +2 kallisto/$gene/abundance.tsv | sort -t $'\t' -k5,5nr | head -5 | cut -f1 | xargs -n 1 samtools faidx "$gene"_transcripts.fasta > "$gene"_TPM_filtered.fasta
      cd $home_dir
      echo
    else
      echo "ERROR: The directory 'igseqr/kallisto/$gene' does not exist."
      exit 1
    fi
  done

###################################### Clean Kallisto Files #####################################

rm -r ./igseqr/kallisto

echo ==============================================================================
echo IgSeqR COMPLETE
echo ==============================================================================
echo
```
